# Gene-Set Enrichment with Mathematical Biology

**DOI:** 10.1101/554212

**Authors:** Amy L Cochran, Kenneth Nieser, Daniel B Forger, Sebastian Zöllner, Melvin G McInnis

## Abstract

Gene-set analyses measure the association between a disease of interest and a *set* of genes related to a biological pathway. These analyses often incorporate gene network properties to account for the differential contributions of each gene. Extending this concept further, mathematical models of biology can be leveraged to define gene interactions based on biophysical principles by predicting the effects of genetic perturbations on a particular downstream function. We present a method that combines gene *weights* from model predictions and gene *ranks* from genome-wide association studies into a weighted gene-set test. Using publicly-available summary data from the Psychiatric Genetics Consortium (n=41,653; ~9) million SNPs), we examine an *a priori* hypothesis that intracellular calcium ion concentrations contribute to bipolar disorder. In this case study, we are able to strengthen inferences from a *P*-value of 0.081 to 1.7×10^−4^ by moving from a general calcium signaling pathway to a specific model-predicted function.

## 1 Introduction

Genetic contributions to disease can be complex and might involve the coordination of a collection of genetic variants in the disruption of one or many biologic pathways. Previous studies of psychiatric conditions provide evidence that a single genetic variant often confers little disease risk despite high heritability [8, 59, 60]. Rather, psychiatric disorders can be *polygenic* [16] — hundreds to thousands of genes of very small effect contribute to the disorder. Genetic risk for an individual is commonly measured by aggregating information from multiple genes into a *polygenic* risk score [2, 23, 29, 44, 52]. Each of these variants might play a small role in the disruption of a pathway, but collectively lead to the development of disease. For this reason, it can be challenging to uncover genetic influences on psychiatric disorders [62]. Computational approaches are emerging to better prioritize candidate genes [3, 6, 11, 12, 17, 21, 26, 27, 32, 39, 46, 65].

Gene-set analyses are a common tool for measuring the association between a disorder and a *set* of genes rather than a single gene [5, 18, 30, 33, 49, 58]. Many statistical tests and software are available to perform gene-set analysis (cf. [33, 45]) to determine whether genes in a particular gene set are significantly associated with a phenotype (self-contained) or whether a phenotype is more strongly associated with genes in a set than genes not in the set (competitive) [18, 33, 64]. Often gene sets are defined based on genes that contribute to a particular biological pathway, which enables identification of pathways that are important for a disorder. This approach likely leads to stronger, more reproducible findings if abnormal pathways are what ultimately contributes to genetic risk [14, 33].

However, biological functions may ultimately drive risk as opposed to an abnormal pathway or single gene variant. Biological functions do not map one-to-one to biological pathways; a function can recruit *some* genes from *multiple* pathways [57]. In bipolar disorder, for example, spontaneous neuronal firing rate differs in stem cells derived from bipolar individuals compared to controls [10, 50]. This cellular function — neuronal firing rate — recruits genes from calcium-mediated signaling (GO:0019722), regulation of action potential (GO:0098900), and chemical synaptic transmission (GO:0007268), among others. Hence, if perturbed biological functions drive disease risk, jointly testing genes in one pathway which includes genes of little impact and ignores genes in other pathways would result in a less powerful gene-set analysis.

Moreover, some genes or gene products play a larger role in the realization of the biologic function. To account for this, some gene-set analyses incorporate information about the network structure of gene interactions [9, 11, 12, 20, 21, 24, 27]. However, the nature of the connections between genes might also vary; interactions might operate in a dynamic and nonlinear way. Greater specificity can be achieved quickly through detailed mathematical models from math biology, which are driven from bottom-up biophysical principles. Efforts within the field have culminated in ModelDB (https://senselab.med.yale.edu/modeldb/) which hosts over 1000 publicly-available models [41]. Examples include models of the hypothalamic-pituitary-adrenal axis, monoamine systems, and circadian rhythms, among others. Model parameters related to genes can be varied to measure the relative contribution of genes to a specific biological function of interest (e.g., firing rate). Incorporating model predictions into gene-set tests might strengthen the link between genes and disorders.

We present a simple method (GEMB: <u>G</u>ene-set <u>E</u>nrichment with <u>M</u>ath <u>B</u>iology) for measuring the association between a disorder and a biological function, based on model predictions. Our method relies on (i) *ranking genes* in decreasing order of association strength to a disorder and (ii) *assigning weights* to a set of genes to reflect their relative contribution to a specific biological function. We illustrate one approach to assigning weights by using pre-existing models from math biology. Ranks and weights are combined into a test for significance of the association between a biological function (as predicted by a neurobiological model) and a disorder.

To demonstrate the utility of our method, we test the hypothesis that intracellular calcium concentrations contribute to bipolar disorder using a detailed model of intracellular calcium concentrations [4]. Bipolar disorder is a severe and chronic psychiatric disorder [7] with estimated heritability at 85% [42]. Genomewide association studies report several susceptibility loci [56], including a voltage-gated calcium gene [25], which remains among the strongest findings to date. Calcium signaling is an incredibly complex process to model [22] but has been implicated in many human diseases [31], including bipolar disorder.

## 2 Materials and Methods

### 2.1 A weighted gene-set statistic

We assume a general set-up of a competitive gene-set test: individuals are phenotyped and analyzed for expression in *n* genes; each gene is measured for association to the phenotype; and a subset of *m* genes are determined to be of interest (see Fig 1 for an overview). From this set-up, we require only the rank of each gene in decreasing order of association strength to the phenotype; genes that are most strongly associated with the phenotype have the highest rank (i.e, closest to 1) and those that are most weakly associated with the phenotype have the lowest rank (i.e., closest to *n*).

**Figure 1:**
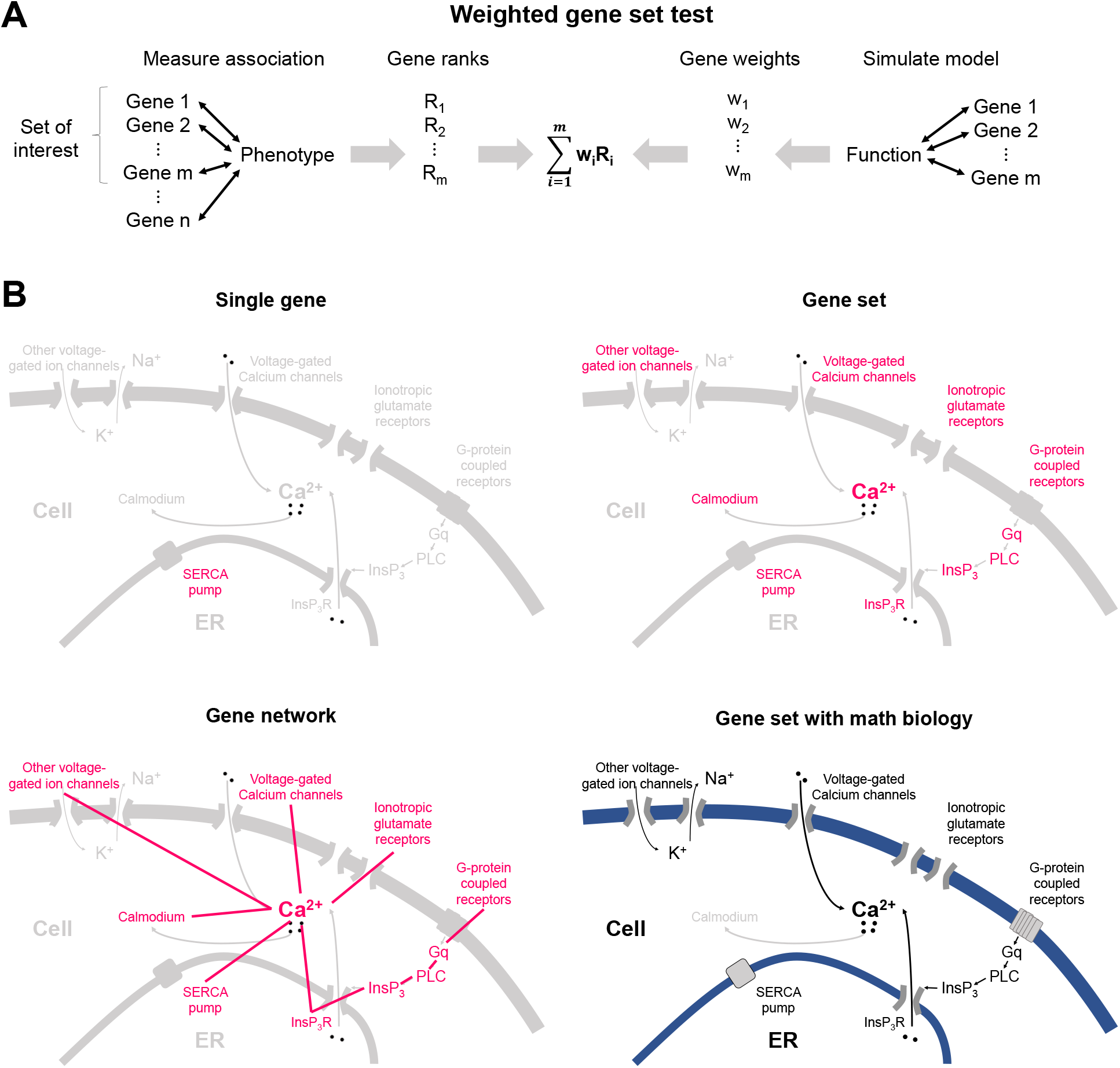
Overview of gene set analysis with math biology. **A)** Genes are ranked based on their association with a phenotype and weighted based on their model-predicted contribution to a specific function. Ranks and genes are combined to perform a weighted gene set test. **B)** Genetic analysis can be performed at the level of either a single gene, a gene set, a gene network, or a gene set connected by math biology. Gene set analysis with math biology uses models to describe connections between genes based on biophysical principles.

We diverge from many gene-set tests by requiring that non-negative weights are assigned to individual genes in the subset of interest. Formally, we require:

- genes labeled 1 to *n*;
- rank *r_i_* ∈ {1, …, *n*} for each gene *i* = 1, …, *n*;
- gene set 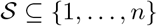; and
- weights *w_i_* ≥ 0 for each gene 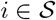 with 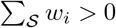.

Without any loss of generality, we assume weights *w_i_* sum to one — we can always re-scale weights so that they sum to one. Then, we define the following test statistic using a weighted sum of the ranks 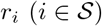:

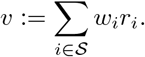

The choice of weights encodes an *a priori* hypothesis about the relative contribution of a gene to the phenotype. As a specific case, we can recover an unweighted gene-set test by setting *w_i_* := 1/*m*. This choice of weights captures the *a priori* hypothesis that each gene in 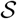 contributes equally to the phenotype (or a lack of support for one gene over another). In this case, the statistic *v* is the average rank of the genes in 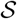. Recalling that a rank of one is assigned to the gene with the strongest association, a value 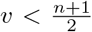 reflects that genes in 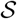 are ranked higher *on average* relative to genes not in 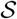. Conversely, a value 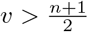 reflects that genes in 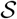 are ranked lower *on average* relative to genes not in 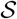. If 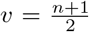, genes in 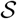 are neither ranked higher nor lower on average relative to genes not in 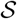. In other words, small *v* suggests an association between the gene set and phenotype. We point out that genes in 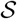 do not need to be evenly distributed in rank to achieve 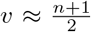; they could be disproportionately ranked close to the average rank 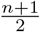 or ranked close to the extreme ranks 1 and *n*.

As another specific case, we could recover a single gene test by setting *w_j_* = 1 for some 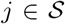 and all other weights to zero. This choice captures the *a priori* hypothesis that gene *i* specifically contributes to the phenotype. The statistic *v* would be the rank of gene *i*. More broadly, setting any weight to zero reflects the hypothesis that the corresponding gene does not contribute to the phenotype. The statistic *v* would be identical in value if we had simply removed the gene from 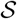. Further, smaller weights means a smaller contribution to *v*.

With more general weights, the statistic *v* is interpreted similarly to the unweighted version, replacing an average of the ranks with a weighted average. Our interpretation is inherited from the fact that *v* – (*n* +1)/2 changes sign when genes are ranked in opposite order and increases when a gene in 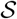 is exchanged for a gene not in 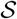 with higher rank. Thus, small *v* can be thought of as genes in 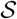 have a higher (weighted) relative rank to genes not in 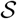.

### 2.2 A weighted gene-set test

To use *v* in a statistical test, we must specify a null distribution. For many gene-set analyses, a common null hypothesis is that the genes in 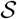 were chosen uniformly at random from the entire set of genes. Under this null hypothesis, we can construct a null distribution for *v* by drawing ranks for genes in our set, *R_i_* for 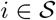, uniformly at random from

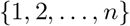

without replacement and calculating

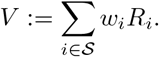

The distribution of the random variable *V* serves as the null distribution for *v*.

The alternative hypothesis is that genes in 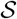 were not chosen uniformly at random. In broad terms, they were chosen because of their relationship to the phenotype. Hence, we are interested in how often *V* with gene ranks chosen randomly suggests a stronger association between set 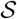 and a phenotype than the statistic *v* determined by the actual association to the phenotype. In other words, we use the probability, or *p*-value, associated with a one-sided test given by

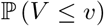

to determine whether *v* is significant. Note, a two-sided test could also be defined by using

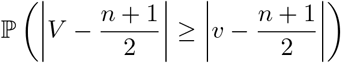

A simple way to estimate 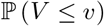 is to use Monte Carlo simulation, where *V* is repeatedly sampled from its distribution and we count how often a sample of *V* is less than or equal to *v*. This computation benefits from the fact that *V* is simple to calculate and can be sampled in parallel. The law of large numbers ensures a Monte Carlo estimate of 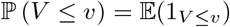 is unbiased and has variance Pr(*V* ≤ *v*)/*k* where *k* is the number of Monte Carlo samples. Alternatively, we could estimate 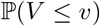 with

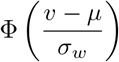

where Φ is a standard normal distribution and 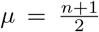 and 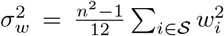. This approximation follows by making the simplifying assumption that ranks *R_i_* are drawn uniformly at random from {1, …, *n*} *with* replacement (as opposed to without replacement) and then noting that the resulting *V* is a sum of independent random variables with respective means 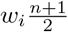 and variances 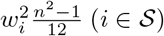. Table 1 compares Monte Carlo estimates of one-sided *P*-values to estimates using a normal approximation.

**Table 1:**
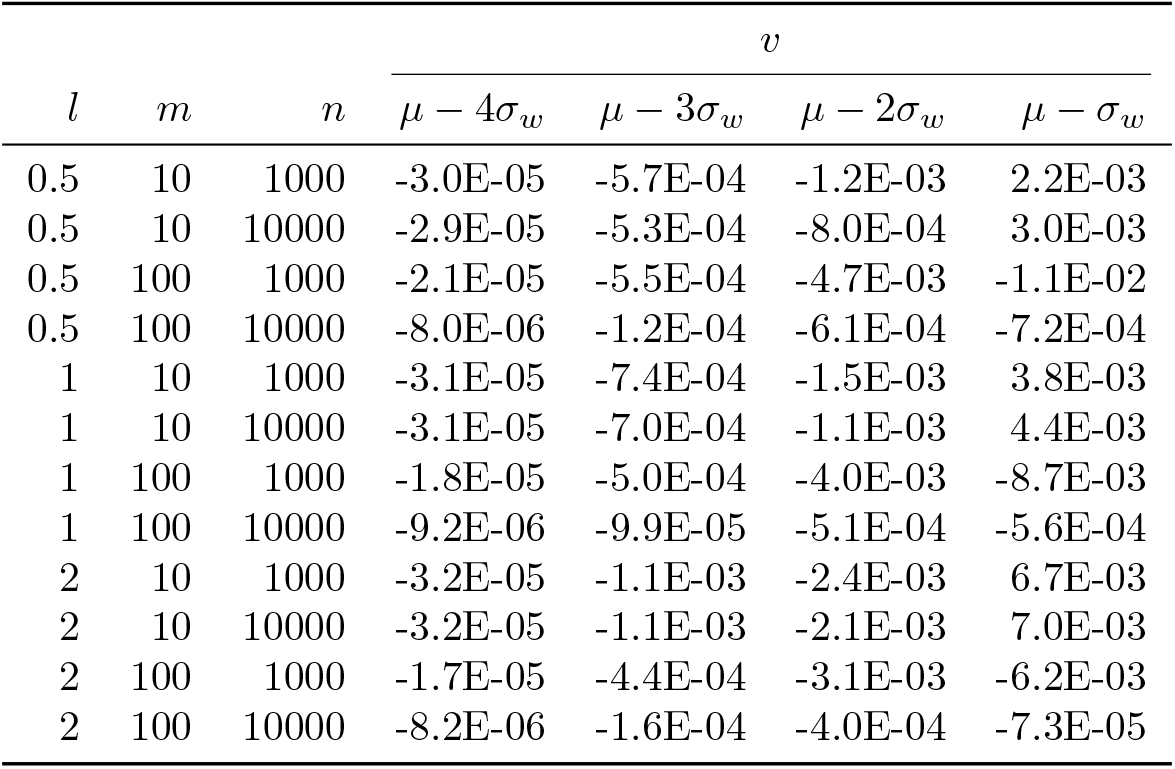
Difference between Monte Carlo estimates of a one-sided *P*-value 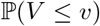 for the weighted gene-set test and estimates using a normal approximation. Weights were defined as *w_i_* ∝ *i^l^* (*i* = 1, …, *m*) for various *l*, assuming 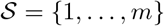. A total of 10^7^ Monte Carlo samples were used in each case.

#### Type I error and Power

Type I error is controlled by the distribution of gene ranks under the null hypothesis of no association between the gene set and the phenotype. Our weighted gene-set test uses the null distribution that arises when any permutation of gene ranks is equally likely. However, the true distribution of gene ranks when there is no association is not clearly defined due to the complex correlations that might exist among genes. Moreover, the null distribution of gene ranks is determined by the method used to generate gene ranks (see [19] for a comparison). It is thus important to choose a method for ranking genes that properly controls Type I error.

Power can be improved with a weighted gene-set test over gene-set or single gene analyses when multiple genes have differential contribution to disease risk. To illustrate, consider *n* genes and a set of two independent genes with very small association to the disease. Under our null hypothesis, gene ranks divided by *n* are approximately uniformly distributed between 0 and 1. A single gene test could assess whether or not each gene’s normalized rank is below some critical value (gray region; Fig 2A). By contrast, a gene-set test could assess whether or not the sum of the two genes’ normalized ranks is below some threshold (blue region; Fig 2A) and a weighted gene-set test could assess whether or not a weighted sum of the two genes’ normalized ranks is below some threshold (green region; Fig 2A). In each case, Type I error is controlled at 0.05 when the rejection region has an area of 0.05.

**Figure 2:**
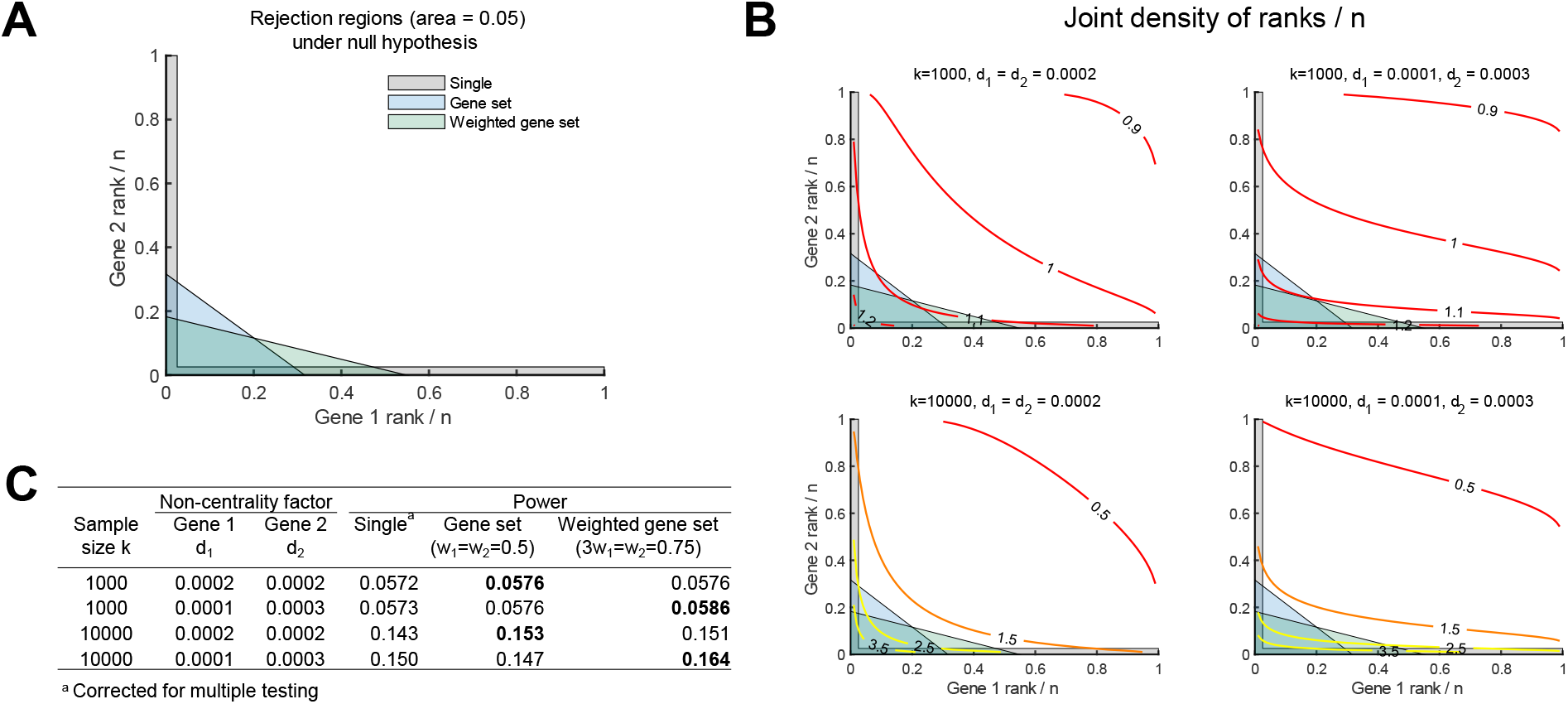
Statistical power. **A)** Possible regions to reject null hypothesis for a single gene test (corrected for multiple testing), a gene set test, and a weighted gene set test. **B)** Joint density functions of ranks divided by n for different sample sizes (*n*) when genes have very small effect sizes (*d*_1_ and *d*_2_). **C)** Statistical power estimated for each case in **(B)**.

To estimate statistical power, we consider a situation when two independent genes of interest are ranked based on an F-test examining if *ν* coefficients are zero when regressing phenotype on gene variables, as is done in MAGMA with *ν* being the number of gene-level principal components used in the regression model [18]. For simplicity, we set *ν* = 10 and assume that P-value of this F-test would be their ranks normalized by the number of genes *n*. For a sample size of *k*, the test statistic for gene 1 and 2 would follow a F-distribution with *ν* – 1 and k-*ν* degrees of freedom under the null hypothesis (no gene–phenotype association). For an alternative distribution, we assume that the test statistic follows a *non-central* F-distribution with *ν* – 1 and *k* – *ν* degrees of freedom and non-centrality parameters *kd*_1_ or *kd*_2_ for gene 1 and 2, respectively. Under this alternative, increasing sample size or non-centrality leads to larger joint densities for normalized ranks near the axes (Fig 2B). Thus with just 2 genes, these changes can improve statistical power, i.e. the probability of arriving at normalized ranks that lie in each reject region (Fig 2C). We expect that this improvement would continue to hold or grow with increases in gene set size and increasingly differential effect sizes. Hence, this example provides support that weighting normalized gene ranks can further increase statistical power by accounting for differential contributions of genes to a disease.

### 2.3 Determining gene weights

Weights can capture any *a priori* hypothesis whether justified by functional data, literature surveys, or experiments. Our goal, however, is more specific: we want weights to reflect the relative contribution of genes to a specific biological function. If we have reason to think certain genes play a large role in the biological function of interest, we upweight them. If genes do not affect the biological function of interest, we downweight them. In this way, our weighted gene-set test incorporates the hypothesis that a specific biological function (captured by the weights) is important to a phenotype.

To inform the choice of weights, we propose a general approach using models from math biology. We start with a neurobiological model that can return a scalar measure of the function of interest. As noted earlier, many models are publicly available through sources such as modelDB. Next, we consult gene databases to identify genes related to one or more model parameters and create a mapping of genes to model parameters. Then, we perform a global sensitivity analysis to measure the relative contribution of each parameter to a specific function of interest. We opt for a global sensitivity analysis based on the partial rank correlation coefficient (PRCC) [40] due to its simplicity. Last, we assign weights to each gene based on the contributions of the model parameter to which it is mapped.

We remark that the association between genes and parameters need not be one-to-one. On one hand, models might not be sufficiently detailed to capture the individual contribution of each gene, so multiple genes may be associated with a single parameter. For example, four genes are known to modulate formation of L-type calcium ion channels, but most mathematical models with L-type ion channels do not include individual parameters to capture the differential contributions of each gene. On the other hand, multiple parameters might be associated with a single gene. For example, models of neuronal action potential often distinguish between sodium currents and persistent sodium currents [51] even though both currents may be regulated by the same gene [37]. We describe how we handled these issues in the context of our case study.

## 3 Results

To illustrate our method, we explore the hypothesis that intracellular calcium ion (Ca^2+^) concentrations in excitable neurons contribute to bipolar disorder. Calcium signaling has been both implicated in bipolar disorder and extensively modeled. Furthermore, this hypothesis was initially tested using our method with a relatively small dataset (n=544) from the Prechter Bipolar Cohort [43] (details in Appendix). Thus, the results reported here represent a replication of our initial finding and validation of an *a priori* hypothesis with a much larger dataset.

### 3.1 Gene ranks

Summary genetic data was obtained on subjects with bipolar disorder (n=20,129) and controls (n=21,524) from the Psychiatric Genomics Consortium (PGC) [53, 54]. Association was measured between 8,958,989 SNPs and bipolar disorder, resulting in *P*-values for each SNP. Data collection and analysis are detailed in Ruderfer et al [54]. Using SNP-level summary data, gene-level association with bipolar disorder was measured using MAGMA software (https://ctg.cncr.nl/software/magma) [18]. Gene locations were defined using NCBI Build 37 (hg19). A total of 3,554,879 (39.68%) SNPs mapped to at least one gene, whereas 18,309 genes (out of 19,427 genes) mapped to at least one SNP. Linkage disequilibrium between SNPs was estimated by MAGMA using reference data files created from Phase 3 of 1,000 Genomes [15]. The set of 18,309 genes were ranked based on their measured association (*P*-value) with bipolar disorder; the smallest *P*-values were ranked closest to 1.

### 3.2 Genes weights

We used a detailed model of an intracellular Ca^2+^ concentration in a hippocampus CA1 pyramidal cell developed by Ashhad and Narayanan in [4]. The model is publicly-available in modelDB (Model 150551) and written with free Neuron software. Furthermore, it captures key contributors to intracellular Ca^2+^ concentrations, including ion transport (K^+^, Na^+^, and Ca^2+^) across the cell membrane; transport of Ca^2+^ into and out of the sarcoplasmic endoplasmic reticulum; synaptic plasticity; and mediating receptors such as inositol triphosphate (InsP_3_), ionotropic glutamate receptors, and metabotropic glutamate recptors (mGR). Finally, the model uses a morphologically realistic three-dimensional representation of a hippocampus CA1 pyramidal cell accompanied by spatial dynamics giving rise to Ca^2+^ waves.

To identify genes of interest, we started with 182 genes making up the *Calcium signaling pathway* (Pathway ko04020) in the Kyoto Encyclopedia of Genes and Genomes (KEGG) [34–36]. Each gene was evaluated for whether it could modulate intracellular Ca^2+^ concentrations in the model using the Gene database from the National Center for Biotechnology Information (https://www.ncbi.nlm.nih.gov/gene). A total of 38 genes could modulate intracellular Ca^2+^ concentrations in the model, by way of ion channels, ion pumps, or receptors. We found three ion channels (Na^+^, A-type K^+^, and delayed rectifying K^+^) and two receptors (NMDA and AMPA) that could affect intracellular Ca^2+^ concentration in the model but had not been associated with genes in the KEGG Calcium signaling pathway. An additional 31 genes were found related to these channels or receptors. Of the 69 genes identified, 4 genes (ATP2B3,CACNA1F,GRIA3,KCND1) were excluded, because they were not associated with gene ranks (described below). A total of 65 genes were analyzed.

For each gene, we identified a parameter that could modulate (up and down) the modeling component related to the gene. For example, channel conductance was associated with ion channel genes. Default parameter values were taken from the simulation in Figure 6 of [4]. Other genes, associated parameters, and default values are summarized in Table 2.

**Table 2:** Calcium genes and associated model parameters. Calcium genes impact either ion channels, ion pumps, or receptors in the Ashhad and Narayanan model [4]. Baseline parameter values were taken from [4].

| Genes | Value | Parameter |
| --- | --- | --- |
| ATP2B[1-2,4] | $0.008 \mu\text{M ms}^{-1}$ | Average rate $\gamma_0$ of $\text{Ca}^{2+}$ flux density |
| ATP2A[1-3] | $0.1 \mu\text{M ms}^{-1}$ | Amplitude $V_{\max}$ of SERCA pump uptake |
| CACNA1[C-D,S] | $0.316 \text{ mS cm}^{-1}$ | L-Type $\text{Ca}^{2+}$ channel conductance $g_{\text{CaL}}$ |
| CACNA1[G-I] | $0.1 \text{ mS cm}^{-1}$ | T-Type $\text{Ca}^{2+}$ channel conductance $g_{\text{CaT}}$ |
| GRM[1,5] | $0.3\text{e-}3$ | Metabotropic glutamate receptor density $[mGR]_0$ |
| GNA[Q,11,14-15] | $100 \text{ ms}^{-1}$ | $G_\alpha$-bound activated PLC formation rate $k_7$ |
| PLC[B1-B4,D1,D3-D4,E1,G1-G2,Z1] | $0.83 \text{ ms}^{-1}$ | PLC$_\alpha$-bound PIP$_2$ formation rate $k_9$ |
| ITPR[1-3] | 1.85 | IP3 receptor density $\bar{g}_{\text{InsP}_3\text{R}}$ |
| GRIN[1,2A-2D,3A-3B] | $1.938107025 \text{ nM s}^{-1}$ | Maximum NMDA receptor permeability $\bar{P}_{\text{NMDA}}$ |
| GRIA[1-2,4] | $1.29207135 \text{ nM s}^{-1}$ | Maximum AMPA receptor permeability $\bar{P}_{\text{AMPA}}$ |
| KCN[A4,C3-C4,D2-D3] | $22 \text{ mS cm}^{-2}$ | A-type $\text{K}^+$ channel conductance $g_{\text{KA}}$ |
| KCN[A1-A3,A6-A7,B1-B2,C1-C2] | $3 \text{ mS cm}^{-2}$ | Delayed rectifying $\text{K}^+$ channel conductance $\bar{g}_{\text{KDR}}$ |
| SCN[1-5,8-11]A | $90 \text{ mS cm}^{-2}$ | $\text{Na}^+$ channel conductance $\bar{g}_{\text{Na}}$ |

With parameters and genes identified, we used the Ashhad and Narayanan model to simulate intracellular Ca^2+^ concentrations during an established protocol for inducing synaptic plasticity at a synapse, namely 900 pulse stimulation at 10 Hz; see Figure 6 in [4]. We simulated 320 samples of parameter sets using Latin-hypercube sampling from a normal distribution with mean given by the respective baseline parameter in [4], standard deviation given by 5% of the respective baseline parameter, and zero correlation. For each parameter set, we simulated intracellular Ca^2+^ and measured average intracellular Ca^2+^ concentrations during initial transients induced in the first three seconds of the simulation.

We estimated the PRCC between each parameter and the measured concentrations controlling for the remaining parameters (Fig 3). We found, for example, a strong positive partial correlation between average intracellular Ca^2+^ concentrations and maximum permeability 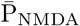 of NMDA receptors and a strong negative partial correlation between average intracellular Ca^2+^ concentrations and the amplitude V_max_ of SERCA pump uptake.

**Figure 3:**
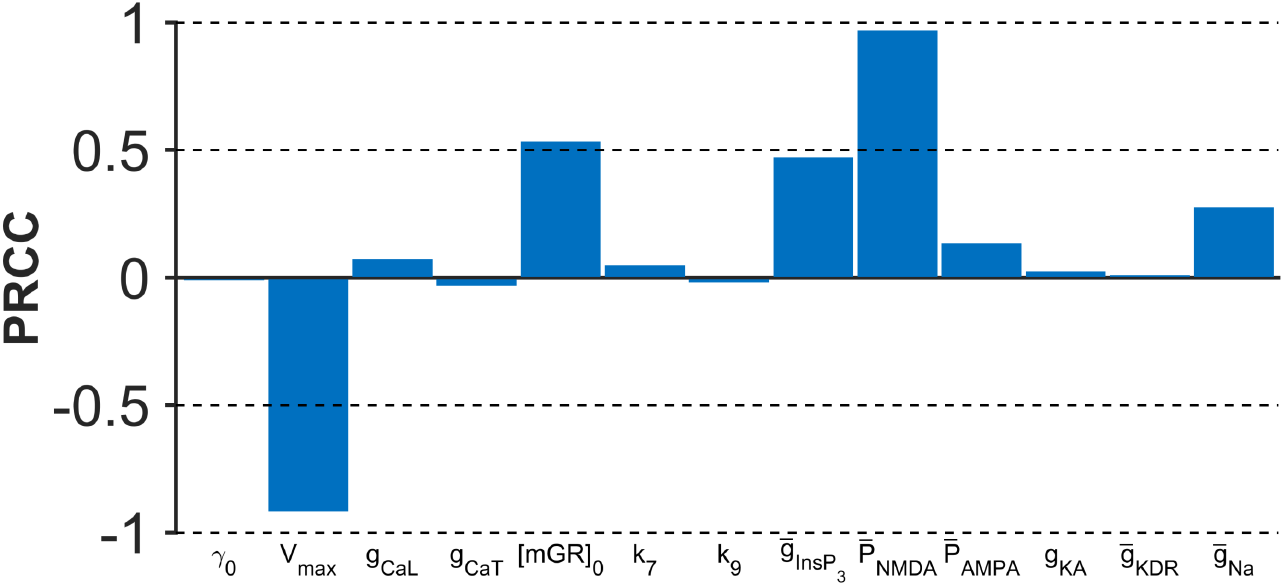
Partial rank correlation coefficient (PRCC) estimated for 13 parameters that modulate intracellular Ca^2+^ concentrations in the Ashhad and Narayanan model [4]. PRCC measures partial correlation between a parameter and the functional measure of interest (average intracellular Ca^2+^ concentration) controlling for the contribution of other parameters.

Based on estimated PRCCs, we defined weights for the 65 genes as follows. For each of the *N_k_* genes assigned to parameter *k* with PRCC *ρ_k_*, we assigned weights |*ρ_k_*|/*N_k_*. We then re-normalized weights to sum to one. Note that we could use any function of *ρ_k_* to assign weights to associated genes. We use only the magnitude of PRCC, since measured associations between genes and phenotypes are not sufficiently specific to reflect the direction of association in addition to the magnitude. We divide by the number of genes assigned to parameter *k*, so that a single component in the model is not weighted heavily simply because there are a large number of genes assigned to the component.

### 3.3 Weighted gene-set test

Combining gene ranks obtained from the genetic analysis with gene weights obtained from the model of calcium signaling, we performed the weighted gene-set test. For comparison, we performed an unweighted gene-set test using all 182 genes from the KEGG Calcium signaling pathway [34–36] by assigning equal weights to all 182 genes. In addition, we performed a typical over-representation analysis with the set of 182 genes. Genes were labeled as significant or not based on a significance level of 0.1 adjusted for false discovery rate [63] (a significance level 0.0044 for our problem); a one-sided Fisher’s exact test was performed to test for over-representation of significant genes in the KEGG calcium signaling pathway compared to genes not in the KEGG calcium signaling pathway. Lastly, since motivation for studying calcium signaling was driven in part by prior PGC results that implicate CACNA1C, we performed the weighted gene-set test without the CACNA1C gene in order to investigate the degree to which our test was driven by the CACNA1C gene.

Our gene-set test (GEMB) showed strong support for our hypothesis that intracellular Ca^2+^ concentration contributes to bipolar disorder (*P*=1.7×10^−4^; Fig 4). The CANA1C gene alone was significant, ranking 10th out of 18,195 genes (*P*=10/18195=5.5× 10^−4^). However, our gene-set test continued to provide strong support for this hypothesis even with the CACNA1C gene removed (*P*=1.9×10^−4^). Thus, our result is only partly driven by the CACNA1C gene. Further, focusing on the entire KEGG Calcium signaling pathway without consideration of differential contributions to biological function provided little support for the hypothesis that calcium signaling is important to bipolar disorder (*P*=0.26 using our method GEMB with equal weights and *P*=0.081 using a one-sided Fisher’s exact test). These discrepancies in *P*-values illustrate how incorporating weights to provide functional specificity could help illuminate biological factors that contribute to a psychiatric disorder.

**Figure 4:**
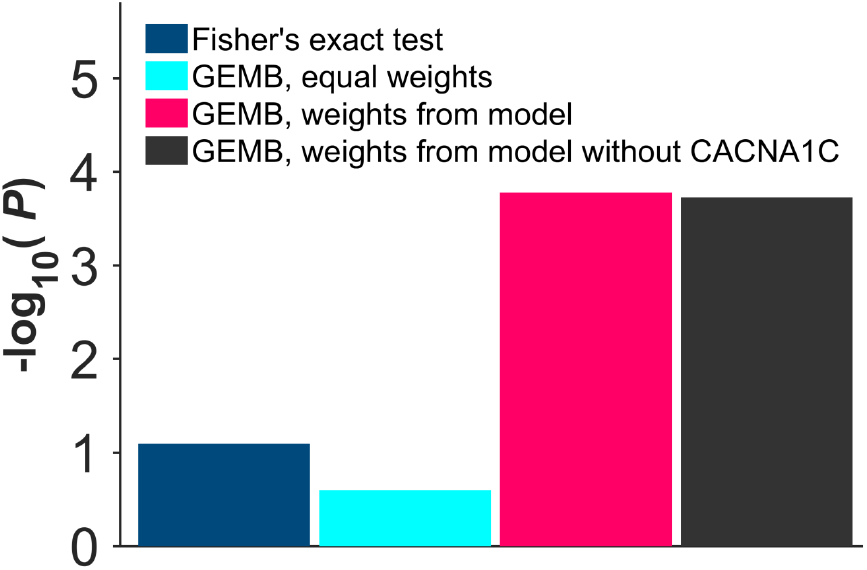
One-sided *P*-values estimated for gene-set tests. Four tests were performed: (1) over-representation test (Fisher’s exact test) applied to the KEGG Calcium signaling pathway; (2) our gene-set test (GEMB) with equal weights applied to the entire KEGG Calcium signaling pathway, (3) our gene-set test (GEMB) with genes related to the Ashhad and Narayanan model [4] and weighted according to their relative contribution to our functional measure of interest (average intracellular Ca^2+^ concentration in an excitable cell); and (4) our gene-set test (GEMB) with weights and genes as determined for (3) but with the CACNA1C gene removed.

## 4 Discussion

We presented a method for examining associations between biological functions and psychiatric disorders which we call GEMB (<u>G</u>ene-set <u>E</u>nrichment with <u>M</u>ath <u>B</u>iology). Central to our method are gene weights that measure the relative contribution of a gene to a particular biological function, which we determine using a neurobiological model. We applied our approach to assess the hypothesis that intracellular calcium ion concentrations are important to bipolar disorder. Gene weights were based on their relative contribution to intracellular calcium ion concentrations, determined by a detailed model of calcium signaling from Ashhad and Narayanan [4]. Gene ranks were obtained using summary genetic data from the Psychiatric Genetics Consortium on bipolar disorder [54], consisting of 20,129 individuals with bipolar disorder and 21,524 controls. Combining gene ranks and weights with our weighted gene-set test, we found strong support for the hypothesis that intracellular calcium concentrations contribute to bipolar disorder (*P*=1.7 × 10^−4^) compared to little support for KEGG Calcium signaling (*P*= 0.081). This result suggests that gene sets defined based on biological pathways may be too broad to capture the genetic effect on a biological *function* that is important to a disorder.

A practical benefit of our weighted gene-set test is that only gene ranks are needed from genetic data. Gene ranks can be shared across researchers more easily and require fewer regulatory and computational resources to analyze compared to full genetic data. Sharing genetic resources and data has become the norm in genetic research, as the community moves towards large consortiums to achieve the sample sizes, level of evidence, and study consistency that are expected. The Psychiatric Genomics Consortium, for example, has “-300 investigators and >75,000 subjects [61], and the National Institute of Mental Health (US) has made genetic data available to researchers. For similar reasons, many popular gene-set tests also need only measures of association (e.g., gene rank) rather than full genetic data [48, 58, 64]. Further, gene ranks can be recovered using MAGMA software from summary data (https://ctg.cncr.nl/software/magma) [18].

Once gene ranks are determined, our method then needs only gene weights from neurobiological models, which too has its benefits. Neurobiological models are numerous, experimentally validated, and publicly available in ModelDB (https://senselab.med.yale.edu/modeldb/). For example, we were able to quickly explore calcium signaling in bipolar disorder, due to the accessibility of a detailed model developed by Ashhad and Narayanan [4] (Model 150551). Similar quick explorations could be used to examine other potentially important biological functions. In searching key words in ModelDB, we found 171 models that contain the concept of *Synaptic Plasticity*, 168 models that contain the concept of *Calcium dynamics*, 47 models that contain the dopamine neurotransmitter, and 9 models that contain the concept of *Circadian Rhythms*, to name a few. Together, these models could annotate genes based on model-predicted functional measures to add to current resources that annotate genes based on biological pathways, such as KEGG [34–36].

With GEMB, neurobiological models may inform genetic studies, but the reverse may also be true: genetic studies may inform neurobiological models. In psychiatry, for instance, there is growing emphasis on team science, affording many opportunities for researchers from the mathematical sciences to help tackle problems [1]. However, just as it is difficult to pin down genes to study in psychiatric disorders, it is also difficult to pin down specific biological processes to study, since abnormal function is found for many neural systems in a psychiatric disorder [28]. Thus, GEMB could help identify, or ground, candidate neurobiological models for studying in psychiatry. The model of Ashhad and Narayanan [4] provides one such example.

Interest is high for ways to incorporate more functional information into gene-set analysis. Network-based approaches, for example, try to incorporate measures of gene relevance based on where they lie in a network in which genes are nodes and gene interactions are edges [3, 9, 11, 12, 20, 21, 24, 26, 27, 32]. Broadly, genes that are more relevant might be more connected or closer to other genes. Other examples include Bayesian approaches to account for overlap between gene sets [6, 39] or approaches based on gene expression levels [17, 46]. A benefit of our method is that it is sufficiently general, such that weights could also be determined from network analysis, experiments or meta-analysis. Weights need only be non-negative and sum to one.

The presented method GEMB was designed to be simple, which has certain limitations. First, our approach does not account for co-expression of genes unlike other genetic analyses [13, 18, 66]. Genes are known to interact in complex ways. Two gene variants, for instance, may lead to increased risk in a disorder that surpasses the additive risk of each variant alone. Second, we do not account for gene interactions in the neurobiological model. Sobol’s method of global sensitivity analysis [55], for example, could measure relative contribution of parameters and their higher-order interactions. Our weighted gene-set test could be extended to incorporate these interactions. Third, neurobiological models are sure to be imperfect, meaning that gene weights are only *predicted* measures of biological function. This issue is, of course, common to all modeling. The question then is not whether using a model leads to the correct answer, but rather whether using models to favor certain genes would strengthen inferences compared to treating the genes equally. This is an empirical question that only continued analyses and applications can answer.

In summary, we have proposed an approach to gene-set analysis that can incorporate a biological hypothesis by weighting genes according to their relative expected contributions. The gain in precision can improve statistical power and strengthen inferences. Most importantly, this method of gene-set analysis can facilitate meaningful biological interpretations which are ultimately necessary in our understanding of the genetic basis of disease.

## Competing Interests

MGM has consulted with and/or received grant funding from Janssen Pharmaceuticals and Takeda Pharmaceuticals; he is a co-owner in Priori-AI, LLC. DBF is the CSO of Arcascope and has equity in the company. Arcascope did not sponsor this research. All other author(s) declare that they have no competing interests.

## A Appendix

### A.1 Coming up with an *a priori* hypothesis

Prior to applying our method to the PCG dataset discussed in the main text, we applied our method to genetic data obtained from the Prechter Bipolar Cohort, a longitudinal cohort of 1,111 individuals [43]. The University of Michigan’s Biomedical Institutional Review Board approved all recruitment, assessment and research procedures (HUM606). Patients provided written informed consent after receiving a complete description of the study. We focused on individuals with bipolar I disorder. Diagnoses of psychiatric illness (e.g. bipolar disorder type I) or lack of psychiatric illness (i.e. control) were determined using the Diagnostic Instrument for Genetic Studies, commonly used in psychiatric research [47]. Diagnoses obtained from the DIGS adhered to DSM-IV diagnostic criteria and were confirmed and re-confirmed annually through a consensus of three clinicians resulting in “best estimate” diagnoses. Participants provided whole blood samples at study intake for genetic testing of specific single nucleotide polymorphisms (SNPs). Methods pertaining to genetic testing are described in detail elsewhere [38]. About 0.5 million SNPs were analyzed initially, which were then used to impute alleles for other SNPs resulting in over 9.8 million SNPs in total.

For the application of our method, we used the same set of genes and the same gene weights obtained from simulation of the Ashhad and Narayanan model [4]. Gene ranks were obtained starting with 428 individuals with BPI and 193 controls without a psychiatric diagnoses. Genetic variation was first analyzed using PLINK software (http://zzz.bwh.harvard.edu/plink/) to account for population stratification and outliers. We performed principal component analysis on SNP data and visualized the participant loadings associated with the first two principal components. We removed any individuals who could be separated from the main cluster in this two-dimensional space either visually or with k-means clustering. This analysis was repeated until there were no participants that could be separated, leaving a total of 377 participants with BPI and 167 controls. Gene-level association to BPI was measured using MAGMA software (https://ctg.cncr.nl/software/magma) [18]. The 10 leading principal components obtained from the final principal component analysis were included as covariates. Genes locations were defined using NCBI Build 38. A total of 18,300 genes were ranked based on the measured association (P-value) with BPI, with smallest P-values ranked closest to 1.

With gene ranks and weights, we performed our weighted gene-set test (GEMB). We again compare our results to an unweighted gene-set test (applying our gene-set test with equal weights) using all 182 genes from the KEGG Calcium signaling pathway [34–36]. We also performed a typical over-representation analysis: genes were labeled as significant or not and then a one-sided Fisher’s exact test was applied to test for over-representation of significant genes in the KEGG Calcium signaling pathway compared to genes not in the KEGG Calcium signaling pathway. However, since the significance level of 0.1 adjusted for false discovery rate yielded no significant genes, we labeled the top 1% of genes as significant [63].

Our gene-set test (GEMB) showed moderate support for our hypothesis that intracellular Ca^2+^ concentration contributes to bipolar I disorder (*P*=0.04). By contrast, focusing on the entire KEGG Calcium signaling pathway provided little support for the hypothesis that Calcium signaling is important to bipolar I (*P*=0.63 using our method GEMB with equal weights and *P*=0.24 using a one-sided Fisher’s exact test). These results provided the impetus to study intracellular calcium concentrations in the larger PCG dataset.

## Notes

#### Summary of Updates

Added a statistical power study; edited writing; added two figures.

https://github.com/cochran4/GEMB

